# Mitogenomic data indicate admixture components of Asian Hun and Srubnaya origin in the Hungarian Conquerors

**DOI:** 10.1101/250688

**Authors:** Endre Neparáczki, Zoltán Maróti, Tibor Kalmár, Klaudia Kocsy, Kitti Maár, Péter Bihari, István Nagy, Erzsébet Fóthi, Ildikó Pap, Ágnes Kustár, György Pálfi, István Raskó, Albert Zink, Tibor Török

**Affiliations:** Department of Genetics, University of Szeged, Hungary; Department of Pediatrics and Pediatric Health Center, University of Szeged, Faculty of Medicine; SeqOmics Biotechnology Ltd. Mórahalom, Hungary; Institute of Biochemistry, Biological Research Centre, Szeged, Hungary; Department of Anthropology, Hungarian Natural History Museum Budapest, Hungary; Department of Biological Anthropology University of Szeged, Hungary; Institute of Genetics, Biological Research Centre, Szeged, Hungary; Institute for Mummies and the Iceman EURAC Bolzano, Italy

**Keywords:** ancient DNA, early Hungarian, next generation sequencing, hybridization capture, Shared Haplogroup Distance, Mitomix

## Abstract

It has been widely accepted that the Finno-Ugric Hungarian language, originated from proto Uralic people, was brought into the Carpathian Basin by the Hungarian Conquerors. From the middle of the 19^th^ century this view prevailed against the deep-rooted Hungarian Hun tradition, maintained in folk memory as well as in Hungarian and foreign written medieval sources, which claimed that Hungarians were kinsfolk of the Huns. In order to shed light on the genetic origin of the Conquerors we sequenced 102 mitogenomes from early Conqueror cemeteries and compared them to sequences of all available databases. We applied novel population genetic algorithms, named Shared Haplogroup Distance and MITOMIX, to reveal past admixture of maternal lineages. Phylogenetic and population genetic analysis indicated that more than one third of the Conqueror maternal lineages were derived from Central-Inner Asia and their most probable ultimate sources were the Asian Huns. The rest of the lineages most likely originated from the Bronze Age Potapovka-Poltavka-Srubnaya cultures of the Pontic-Caspian steppe, which area was part of the later European Hun empire. Our data give support to the Hungarian Hun tradition and provides indirect evidence for the genetic connection between Asian and European Huns. Available data imply that the Conquerors did not have a major contribution to the gene pool of the Carpathian Basin, raising doubts about the Conqueror origin of Hungarian language.

## Introduction

Foundation of the Hungarian state is connected to the conquering Hungarians, which arrived from the Pontic steppes and occupied the Carpathian Basin at 895-905 AD as a confederation of seven tribes under the leadership of prince Árpád. Modern Hungarians are generally identified as descendants of the conquering Hungarians (hence shortened as Conquerors). Until the middle of the 19^th^ century it was generally accepted that Hungarians were kinsfolk of the Huns and Scythians, besides Árpád was a direct descendant of the great Hun leader Attila. Hun-Hungarian affinity was declared in Hungarian and foreign written sources and has been maintained in Hungarian folk memory [1–3]. Most Hungarian medieval chronicles describe the conquer as the “second incoming of the Hungarians” [1]. In the second half of the 19th century the Hungarian language was reclassified as belonging to the Uralic branch of the Finno-Ugric language family [4]. Philological arguments launched a reevaluation of previous assumptions and as a result, the credibility of medieval historical sources, including Hun-Hungarian relations, has been questioned. In following decades the Hungarian Conquerors were deemed descendants of hypothetic proto Uralic people, the putative common ancestors of people belonging to this language family. Lately most philologists proclaim separability of linguistic and genetic relations, but appearance of the Hungarian language in the Carpathian Basin is explicitly linked to the Conquerors [5].

The possible genetic relation of modern Hungarians to Finno-Ugric groups was tested in several studies [6–8], however all these found Hungarians being genetically unrelated to Uralic people. One of the latest studies [9] reported that a Y-chromosome haplogroup (*N-L1034*) is shared between 4% of the Hungarian Seklers (Hungarian-speaking ethnic group living in Transylvania) and 15% of the closest language relatives the Mansis, though the same marker is also present in Central Asian Uzbeks and has been detected just in one Hungarian [10]. These results indicated that Uralic genetic links hardly exist in modern Hungarians.

The genetic composition of the Conquerors was also analyzed in several ancient DNA (aDNA) studies [11–13] and indeed, all these detected significant presence of East Eurasian major mtDNA haplogroups (Hg-s), which are rare in modern Hungarians but are found in Uralic people. Another study [14] showed the presence of *N-Tat* (*M46*) Y-chromosome marker (a major clade of the above mentioned *N-L1034*) in two of the Conqueror samples and one living Sekler, which was interpreted as a Finno-Ugric link. It is notable that in the latest studies [12,13] population genetic analysis also indicated considerable Central Asian affinity of the Conquerors. However all these studies applied low resolution, error prone PCR based aDNA methods [15] and high resolution Next Generation Sequencing (NGS) data from the Conquerors has not been available yet, though power of discrimination of the entire mitochondrial genome over the hypervariable regions (HVR) has been shown to be huge [16].

In order to elicit the genetic origin and relationships of the Conquerors, we set out to assemble a full length mtDNA sequence database from the earliest Conqueror cemeteries. Full length mitogenomes are the most informative source of maternal population histories, as some of the subclades have very distinctive geographic distribution [17,18], reviewed in [19]. Thus the availability of ancient mitogenomes obtained with NGS greatly enhanced the resolution of the phylogeographic approach, making it possible to refine the view of peopling of the Americas [20] and Europe [21–23]. We also made use of this approach by comparing the mtDNA genomes of 102 Conqueror individuals to available public databases. Applying phylogenetic analysis we could allocate the ultimate geographical origin of the Hg lineages to various regions of Eurasia, while population genetic results pointed at more recent source populations in Tuva, Central Asia, today’s Belarus and Volga district, providing new information about the origin of the Conquerors, which is reconcilable with historical sources.

## Materials and Methods

### Archaeological background

In the 10^th^ century a uniform well distinguishable new archaeological culture appeared in the Carpathian Basin which can be connected to the historical record of the conquering Hungarians. We extracted ancient DNA from 102 Conqueror individuals, derived from 8 different cemeteries (Suppl. text). As one of our purposes was to characterize the entire population from a few early Conqueror cemeteries, the majority of samples came from three cemeteries of Karos-Eperjesszög, representing the earliest Conquerors in the Carpathian basin. These three cemeteries are located in the upper Tisza river region on neighboring sand dunes a few 100 meters from each other, with the richest archaeological findings of the period, and were probably used by contemporary neighboring communities from the last years of the ninth century to the middle of the tenth century, based on dating with coins and comparative analysis of archaeological findings [24]. Basic archaeological description of the cemeteries were given in [13], further details are provided in Suppl text and S1 Table.

We have sequenced mitogenomes from all of the available 11 remains of the Karos1 cemetery, 49 out of 69 graves from the Karos2 cemetery, and 18 out of 19 graves from the Karos3 cemetery. For comparison we also sequenced 24 mitogenomes from the following Conqueror cemeteries: 10 individuals from Kenézlő-Fazekaszug1-2, 8 from Sárrétudvari-Hízóföld, two from Szegvár-Oromdűlő, and one from each of the following: Harta-Freifelt, Magyarhomoróg, Orosháza-Görbicstanya and Szabadkígyós-Pálliget (Suppl. text). None of the individuals in the cemeteries showed signs of violent death, most had ceremonial burial according to nomadic traditions and the ratio of children and elderly individuals indicate a vital community (S1 Table).

### NGS sequencing

Details of the aDNA purification, hybridization capture, sequencing and sequence analysis methods are given in [15]. In order to authenticate the results, we considered the latest recommendations of [25] throughout of the experiments. We tried to apply the modifications recommended by [26] on a few samples (Kenézlő-Fazekaszug/ 1027, 1044, 1045 and 10936, Sárrétudvari-Hízóföld/ 66 and 103), but in our hands this method gave rather varying coverage. In some of the samples (Karos2/2, 17, 18, 33, 44, 67, Karos3/7, 9, 11, 13, 17, 18, Sárrétudvari-Hízóföld/9-anc11, Sárrétudvari-H/2, 4, and Kenézlő-F/1025, 1031, 1036, 1041, 1042) we decreased the recommended USER and UGI concentrations of [27] to half (0.03 U/μL) and at the same time increased the incubation time from 30 minutes to 40 minutes. This modification removed uracils with comparable efficiency to the original method.

Details of NGS data are shown in S3 Table. Most genomes had satisfactory coverage, but we also included several low coverage sequences, whose Hg-s could be unmistakably classified, as these revealed meaningful maternal relationships within and between cemeteries. We used the Schmutzi algorithm [28] to estimate contamination and to verify consensus sequences. In most cases Schmutzi measured low contamination levels, however in 12 samples it estimated 99% contamination As in some of these “highly contaminated” samples Schmutzi detected low contamination when it was run with unfiltered BAM files, containing larger number of damaged molecules, we infer that partial UDG treatment in these samples left not enough transitions in key positions for Schmutzi to start with in the first iteration on filtered BAMs. For these samples contamination was estimated traditionally by calculating the proportion of reads which did not correspond to the consensus sequence in diagnostic positions [15]. Sequences were deposited to the European Nucleotide Archive under accession number PRJEB21279.

### Phylogenetic study

We have downloaded all available modern (n=32683) and ancient (n=564) complete mtDNA genome sequences from the NCBI and European Nucleotide Archive databases or requested them from the authors. This database was also augmented with 314 mitogenomes including 272 new Hungarian ones described in [29]. Then we determined the haplogroups of all sequences with the HaploFind program [30], and arranged them according to haplogroups. Next we selected each subset of sequences (28-180/Hg) corresponding to the Hg of individual Conqueror samples. Selected sequence subsets were aligned with MAFFT version 7 [31,32] using progressive G-INS-1 setting. Aligned multifasta groups were converted into Nexus file with MEGA [33], then Median-Joining networks [34] were drawn with PopART [35]. Finally phylogeographic connections were inferred by looking up the geographic origin of the closest matching samples from the literature (S1 Figure).

### Population genetic study

We have created an Eurasian population database by grouping those mtDNA genomes according to their geographic origin, for which this information was available. Our population database contains 12224 modern samples from 62 Eurasian populations (S4, S5 Tables) not considering India and Southeast Asia. In cases when populations were underrepresented we grouped related neighboring groups, like Mansis with Khantys, Belgians with Dutch etc., as listed in S4 Table. We also created a similar mitogenomic population database from 25 ancient Eurasian populations including 495 sequences, though most of these contain low number of samples (S4, S5 Tables).

We compared the genetic similarity of populations with two independent methods. We applied the traditional sequence based method calculating pair-wise population differentiation values (Fst) with Arlequin 3.5.2.2 [36] from entire mtDNA genomes (S7 Table) assuming a Tamura & Nei substitution model (Tamura and Nei, 1993) with a gamma value of 0.325. Significant variations in Fst values were tested by 10,000 permutations between populations. As individual insertions and deletions make the alignment of multiple mtDNA genomes troublesome, only variable positions were aligned, and insertions and deletions were recoded to SNP-s as follows. Whole mtDNA genome fasta files were aligned to the NC_012920 human mtDNA reference sequence by an IUPAC code aware in-house aligner using the Needleman–Wunsch algorithm with weight parameters: match 6, IUPAC2match (R, Y, M, W, S, K) 3, IUPAC3match (B, D, H, V) 2, IUPAC4match (N) 1, mismatch −12, gap open −24, gap extend −6. Modern sequences with more than 500 missing or uncertain nucleotides (nt.) were excluded from further analysis. Then all nt. positions where any variation was detected were outputted to VCF files. Since Arlequin cannot manage VCF files SNPs, deletions and insertions were recoded by the following rules: nt-s with no variation at the given position were coded as the reference nt.; SNPs with variation were coded as the alternate allele; all insertions were coded as additional nt. letters, C for samples with reference sequence and T for samples containing the insertion; all deletions were also coded as additional nt. letters, T for samples with reference sequence and C for samples containing the deletion. Then Arlequin input files (arp) were generated from the recoded DNA sequences.

Multidimensional scaling (MDS) was applied on the matrix of linearized Slatkin Fst values [37] and visualized in the two-dimensional space using the cmdscale function implemented in R 3.0.3 [38].

In a second novel approach we also calculated so called Shared Haplogroup Distance (SHD) values between populations [29]. This approach takes into account solely haplogroup defining SNP-s and ignores all other sequence differences. This method considers that all individuals within the same sub-Hg were descended from a single foremother, therefore their maternal lineages are more closely related to each-other than to individuals of neighboring sub-Hg-s, regardless of other sequence differences. While Fst based calculations are best suited for measuring evolutionary distances between not admixing populations, SHD based distance reveals recent admixtures more accurately [29]. This method calculates a distance value between 0-1, which is minimum between populations containing the same sub-haplogroups with identical frequencies, and maximum between populations with no sub-Hg overlap. We used corrected SHD vales, which also takes into account the mutation and fixation rate on the mtDNA genome, thereby allows some connection between progenitor and progeny Hg lineages [29]. Pair-wise SHD distances were calculated between all 87 ancient and modern populations from the frequency of 1942 sub-Hg-s occurring in any of them (S8 Table).

We have also used another novel algorithm called MITOMIX, which computes all possible combinations and proportions of *K* populations to find the best fitting admixtures with the smallest SHD values from a test population [29]. In our experience most test populations are adequately admixed from 3-6 other populations, as *K* values greater than 6 do not significantly improve the result. Theoretically MITOMIX can accurately reconstruct past population admixtures if representative data are available from all periods and locations, but it allows meaningful insights even from limited data. With this method we have calculated the best population admixtures for the Conquerors, as well as for their possible source populations (S10-S15 Tables).

### Craniofacial Reconstructions

The sculpting craniofacial reconstructions of three skulls from the Karos cemeteries (Figure 6?) were carried out by Gyula Skultéty in cooperation with the Hungarian Natural History Museum, Department of Anthropology [39,40]. During the facial reconstruction, soft tissue layers were grafted back onto the plaster copy of the skulls carefully following the bone conformation to accurately recreate the facial features according to the published guidelines [41–43]. Facial reconstruction was performed by traditional sculpting anatomy, that is plasticine muscles were attached in their anatomically correct position [40,44]. The width of a muscle was determined by the ruggedness of the bone surface by means of a table compiled from measurements taken from 45 different points of the skull. These data have been collected by scientific methods [45].

## Results

### Phylogenetic study

Using the NGS sequencing method combined with target enrichment, we could obtain 102 ancient mitogenome sequences, 78 of which are first reported in this paper, while 24 had been reported in [15]. The 102 sequences belong to 67 sub-Hg-s, and first we elucidated the phylogenetic relations of each Hg-s using M-J Networks as shown in S1 Figure. With a few exceptions M-J networks typically separated neighboring sub-Hg-s to different branches. The closest sequence matches pointed at a well defined geographical region in most cases, which is indicated next to the phylogenetic trees and is summarized on Figure 1. Phylogenetic trees revealed that the Conqueror maternal lineages originated from two distant geographical regions; 31 were unequivocally derived from East Eurasia, while 60 from West Eurasia. The remaining 11 Conqueror Hg-s are ubiquitous in Eurasia. Out of the 60 West Eurasian lineages 13 are characteristic for modern Northwestern Europeans, while 7 have primarily Caucasus-Middle-East distribution.

**Fig. 1.**
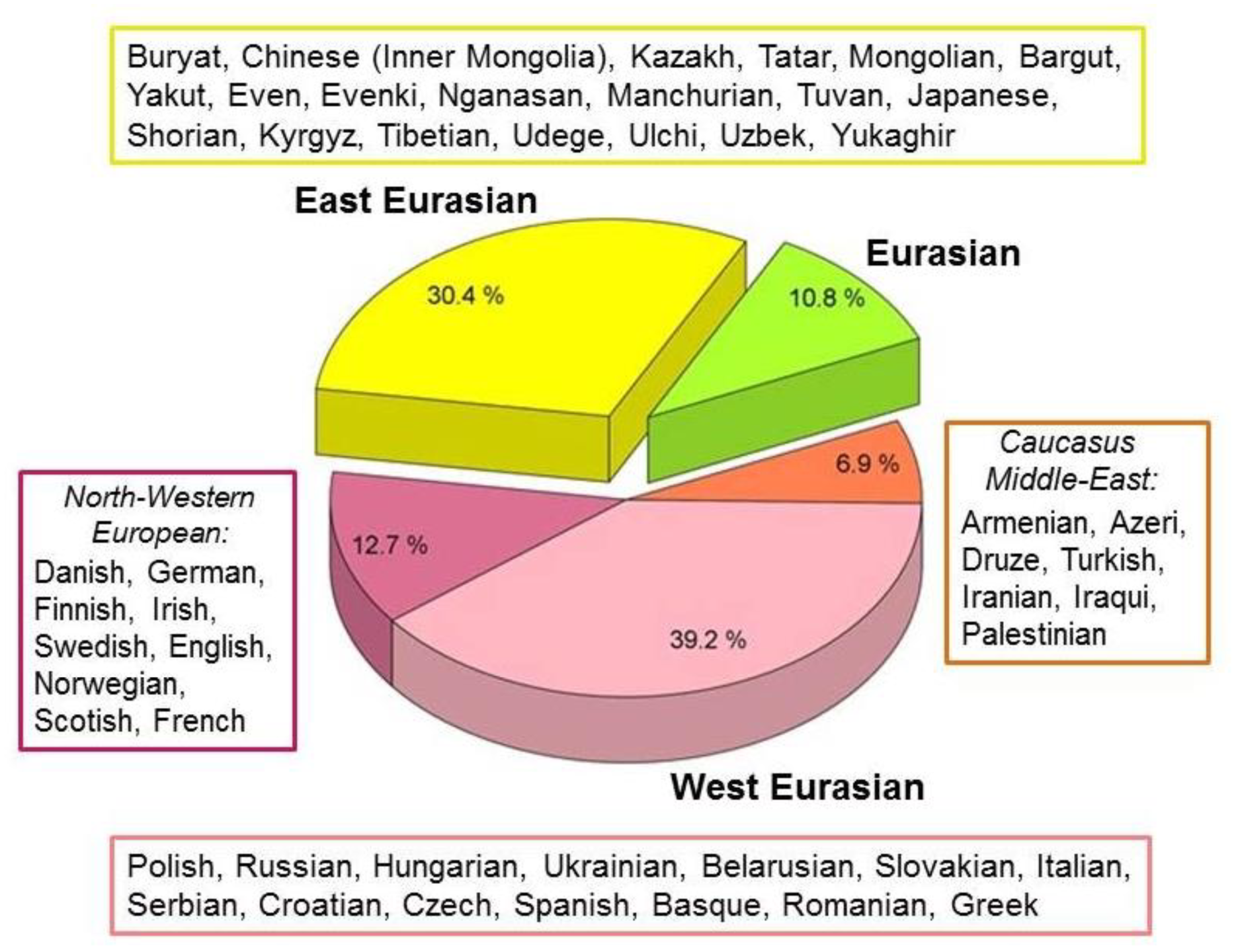
Phylogeographic origin of the 102 Conqueror maternal lineages. Data are summarized from S1 Figure. Origin of modern individuals with closest matches to Conqueror sequences are listed next to the indicated regions, ordered according to frequency of appearances.

Many of the sequences showed close matches with ancient samples (S1 Figure) indicating affinity to ancient cultures, which is summarized on Figure 2. Most lineages contain ancient sequences of steppe origin, the most recurrent being the Bronze Age Srubnaya culture (Timber-grave, EU Steppe LMBA on Figure 2) with 10 closely related Conqueror sequences. The prominent frequency of Hg *N1a1a1a1a*, represented by 7 samples, is a revealing genetic signal for Central Asian origin, as current distribution of this Hg is restricted to Kazakhstan, Altai, Buryat Republic and Russia, attesting that these areas were the center of expansion [46]. This Hg was detected in a Bronze Age Sintashta sample from Kazakhstan [47], an Iron Age Pazyryk Scythian [48] and an early Sarmatian sample [49], while its progenitor Hg *N1a1a1a1* has a wide Eurasian distribution [46]. Our phylogeographic data (S1 Figure, Network 36) imply a probable expansion of *N1a1a1a1* from the European Pontic Steppe to Central Asia around the Bronze Age and its sub-clade *N1a1a1a1a* from Central Asia both to Inner Asia and back to Europe from the Iron Age. Close affinity to Sintastha, Andronovo, Karasuk, Okunevo and Scythian steppe cultures is also revealed by 16 other samples. The European Late Neolithic-Early Bronze Age (EULNBA) is represented by 13 closely related samples, and the Nordic and Baltic Bronze Age affinity appears significant as well. Two *D4j12* individuals (S1 Figure, Network 9) from the Karos1 cemetery had close match with an European Hun sample from the Carpathian Basin [50], indicating possible European Hun affinity of these maternal lineages. The only close match of the Karos2/30 individual carrying *K1f* Hg (S1 Figure, Network 35) was the Iceman, indicating Chalcolithic European origin. In addition Armenian Bronze Age, European Neolithic, Samara Eneolithic, Baltic Combed Ware, Yamnaya and Celtic samples appear on the list. Multiple colors appearing simultaneously on Figure 2 indicate considerable overlap between most of the ancient relations.

**Fig. 2.**
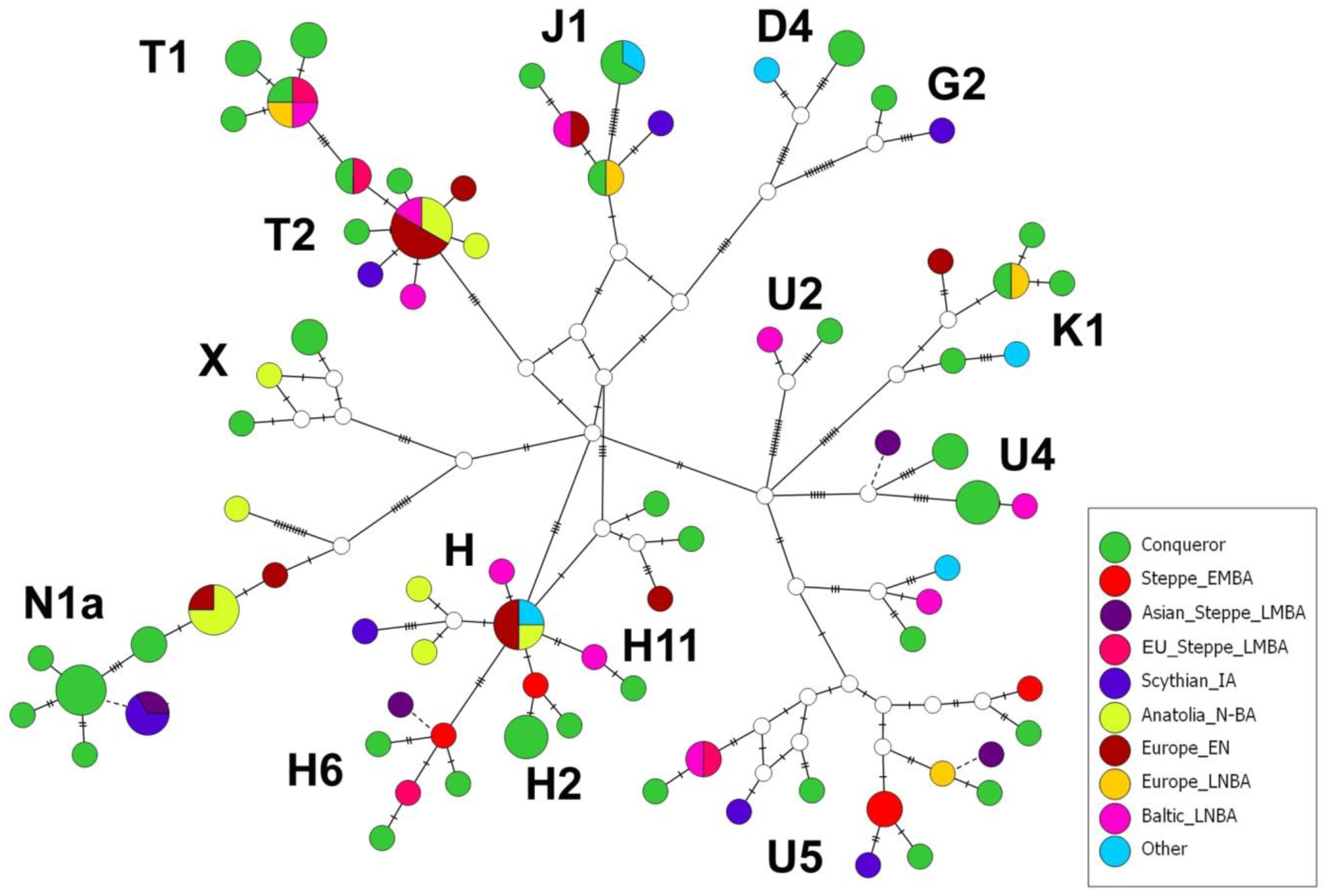
M-J Network drawn from ancient sequences with major Hg-s labeled. Conqueror sequences having closest sequence matches with any of the ancient cultures are portrayed from S1 Figure. Informative incomplete sequences, which could not be properly aligned, are indicated with dashed line. Color coded groups represent the following cultures: Steppe EMBA (Afanasevo, Yamnaya Samara, Eneolith Samara); Asian Steppe LMBA (Sintastha, Andronovo, Okunevo, Karasuk); EU Steppe LMBA (Potapofka, Srubnaya); Scythian IA (Scythian Samara, Early Sarmatian Ural, Pazyryk); Anatolian N-BA (Neolith Turkey, Armenian Neolith-Early Bronze Age-Iron Age); Europe EN (European Early-Mid Neolithic); Europe LNBA (Post LBK Poland, Central LNBA Europe, European Chalcolithic, Bell Baker, Remedello Italy); Baltic LNBA (Narva, Trzciniec, Baltic Corded Ware, Scandinavian Bronze Age), Other (Celtic, Roman, Combed Ware, Holocene Italy, European Hun).

### Population genetic study

The presence of 30% East Eurasian and 60% West Eurasian Hg lineages raises the possibility that the Conquerors may have been assembled from at least two distant source populations. To model all possible scenarios we assembled 5 hypothetic Conqueror subpopulations; a) all 102 sequences together; b) the 60 European lineages; c) the 60 European lineages together with the 11 Eurasian lineages; d) the 31 East Eurasian lineages; e) the 31 East Eurasian lineages together with the 11 Eurasian lineages (S6 Table), and measured their genetic distances from all recent and ancient populations with two independent methods. Fst and SHD distance matrixes are provided in S7-9 Tables, and are summarized in Table 1.

**Table 1.**
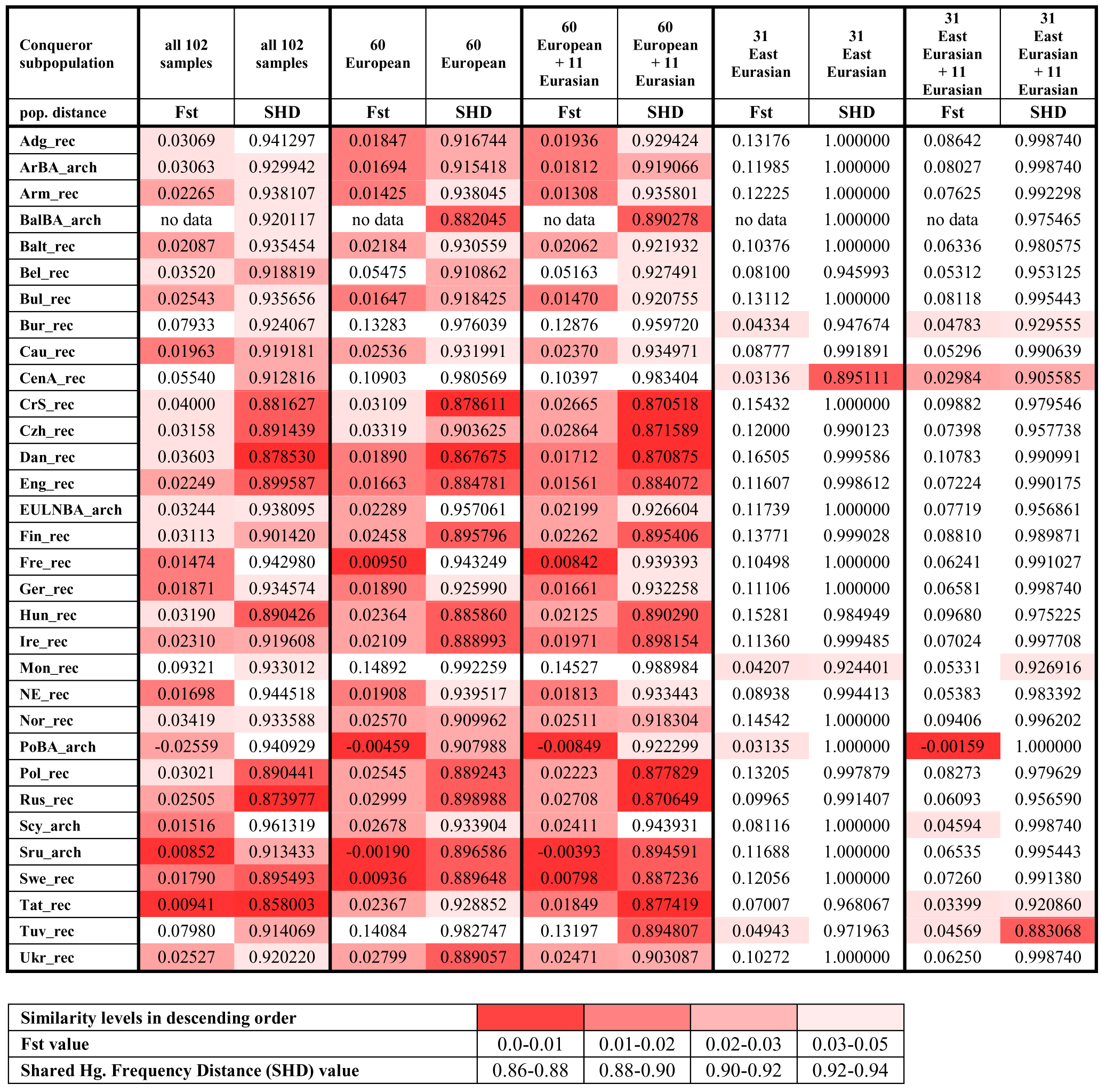
Fst and SHD distances of modern and ancient populations measured from different Conqueror subpopulations. Only populations which showed close distance values with both methods for any of the Conqueror subpopulations are displayed here from S9 Table. Abbreviations of population names are given in S4 Table.

| Conqueror subpopulation | all 102 samples | all 102 samples | 60 European | 60 European | 60 European + 11 Eurasian | 60 European + 11 Eurasian | 31 East Eurasian | 31 East Eurasian | 31 East Eurasian + 11 Eurasian | 31 East Eurasian + 11 Eurasian |
| --- | --- | --- | --- | --- | --- | --- | --- | --- | --- | --- |
| pop. distance | Fst | SHD | Fst | SHD | Fst | SHD | Fst | SHD | Fst | SHD |
| Adg_rec | 0.03069 | 0.941297 | 0.01847 | 0.916744 | 0.01936 | 0.929424 | 0.13176 | 1.000000 | 0.08642 | 0.998740 |
| ArBA_arch | 0.03063 | 0.929942 | 0.01694 | 0.915418 | 0.01812 | 0.919066 | 0.11985 | 1.000000 | 0.08027 | 0.998740 |
| Arm_rec | 0.02265 | 0.938107 | 0.01425 | 0.938045 | 0.01308 | 0.935801 | 0.12225 | 1.000000 | 0.07625 | 0.992298 |
| BalBA_arch | no data | 0.920117 | no data | 0.882045 | no data | 0.890278 | no data | 1.000000 | no data | 0.975465 |
| Balt_rec | 0.02087 | 0.935454 | 0.02184 | 0.930559 | 0.02062 | 0.921932 | 0.10376 | 1.000000 | 0.06336 | 0.980575 |
| Bel_rec | 0.03520 | 0.918819 | 0.05475 | 0.910862 | 0.05163 | 0.927491 | 0.08100 | 0.945993 | 0.05312 | 0.953125 |
| Bul_rec | 0.02543 | 0.935656 | 0.01647 | 0.918425 | 0.01470 | 0.920755 | 0.13112 | 1.000000 | 0.08118 | 0.995443 |
| Bur_rec | 0.07933 | 0.924067 | 0.13283 | 0.976039 | 0.12876 | 0.959720 | 0.04334 | 0.947674 | 0.04783 | 0.929555 |
| Cau_rec | 0.01963 | 0.919181 | 0.02536 | 0.931991 | 0.02370 | 0.934971 | 0.08777 | 0.991891 | 0.05296 | 0.990639 |
| CenA_rec | 0.05540 | 0.912816 | 0.10903 | 0.980569 | 0.10397 | 0.983404 | 0.03136 | 0.895111 | 0.02984 | 0.905585 |
| CrS_rec | 0.04000 | 0.881627 | 0.03109 | 0.878611 | 0.02665 | 0.870518 | 0.15432 | 1.000000 | 0.09882 | 0.979546 |
| Czh_rec | 0.03158 | 0.891439 | 0.03319 | 0.903625 | 0.02864 | 0.871589 | 0.12000 | 0.990123 | 0.07398 | 0.957738 |
| Dan_rec | 0.03603 | 0.878530 | 0.01890 | 0.867675 | 0.01712 | 0.870875 | 0.16505 | 0.999586 | 0.10783 | 0.990991 |
| Eng_rec | 0.02249 | 0.899587 | 0.01663 | 0.884781 | 0.01561 | 0.884072 | 0.11607 | 0.998612 | 0.07224 | 0.990175 |
| EULNBA_arch | 0.03244 | 0.938095 | 0.02289 | 0.957061 | 0.02199 | 0.926604 | 0.11739 | 1.000000 | 0.07719 | 0.956861 |
| Fin_rec | 0.03113 | 0.901420 | 0.02458 | 0.895796 | 0.02262 | 0.895406 | 0.13771 | 0.999028 | 0.08810 | 0.989871 |
| Fre_rec | 0.01474 | 0.942980 | 0.00950 | 0.943249 | 0.00842 | 0.939393 | 0.10498 | 1.000000 | 0.06241 | 0.991027 |
| Ger_rec | 0.01871 | 0.934574 | 0.01890 | 0.925990 | 0.01661 | 0.932258 | 0.11106 | 1.000000 | 0.06581 | 0.998740 |
| Hun_rec | 0.03190 | 0.890426 | 0.02364 | 0.885860 | 0.02125 | 0.890290 | 0.15281 | 0.984949 | 0.09680 | 0.975225 |
| Ire_rec | 0.02310 | 0.919608 | 0.02109 | 0.888993 | 0.01971 | 0.898154 | 0.11360 | 0.999485 | 0.07024 | 0.997708 |
| Mon_rec | 0.09321 | 0.933012 | 0.14892 | 0.992259 | 0.14527 | 0.988984 | 0.04207 | 0.924401 | 0.05331 | 0.926916 |
| NE_rec | 0.01698 | 0.944518 | 0.01908 | 0.939517 | 0.01813 | 0.933443 | 0.08938 | 0.994413 | 0.05383 | 0.983392 |
| Nor_rec | 0.03419 | 0.933588 | 0.02570 | 0.909962 | 0.02511 | 0.918304 | 0.14542 | 1.000000 | 0.09406 | 0.996202 |
| PoBA_arch | -0.02559 | 0.940929 | -0.00459 | 0.907988 | -0.00849 | 0.922299 | 0.03135 | 1.000000 | -0.00159 | 1.000000 |
| Pol_rec | 0.03021 | 0.890441 | 0.02545 | 0.889243 | 0.02223 | 0.877829 | 0.13205 | 0.997879 | 0.08273 | 0.979629 |
| Rus_rec | 0.02505 | 0.873977 | 0.02999 | 0.898988 | 0.02708 | 0.870649 | 0.09965 | 0.991407 | 0.06093 | 0.956590 |
| Scy_arch | 0.01516 | 0.961319 | 0.02678 | 0.933904 | 0.02411 | 0.943931 | 0.08116 | 1.000000 | 0.04594 | 0.998740 |
| Sru_arch | 0.00852 | 0.913433 | -0.00190 | 0.896586 | -0.00393 | 0.894591 | 0.11688 | 1.000000 | 0.06535 | 0.995443 |
| Swe_rec | 0.01790 | 0.895493 | 0.00936 | 0.889648 | 0.00798 | 0.887236 | 0.12056 | 1.000000 | 0.07260 | 0.991380 |
| Tat_rec | 0.00941 | 0.858003 | 0.02367 | 0.928852 | 0.01849 | 0.877419 | 0.07007 | 0.968067 | 0.03399 | 0.920860 |
| Tuv_rec | 0.07980 | 0.914069 | 0.14084 | 0.982747 | 0.13197 | 0.894807 | 0.04943 | 0.971963 | 0.04569 | 0.883068 |
| Ukr_rec | 0.02527 | 0.920220 | 0.02799 | 0.889057 | 0.02471 | 0.903087 | 0.10272 | 1.000000 | 0.06250 | 0.998740 |

**Table 1. Fst and SHD distances of modern and ancient populations measured from different Conqueror subpopulations.**
| Similarity levels in descending order |  |  |  |  |
| --- | --- | --- | --- | --- |
| Fst value | 0.0-0.01 | 0.01-0.02 | 0.02-0.03 | 0.03-0.05 |
| Shared Hg. Frequency Distance (SHD) value | 0.86-0.88 | 0.88-0.90 | 0.90-0.92 | 0.92-0.94 |

The Fst and SHD methods gave comparable results, close distances measured with either method mostly appear close with the other method too (S9 Table). Exceptions well illuminate the differences between the two approaches; SHD is more susceptible to detect admixture [29], as it reveals the presence of Buryat (Bur), Central Asian (CenA), Mongolian (Mon) and Tuvinian (Tuv) components in the entire Conqueror population, which are only detected in the East Eurasian subset by Fst (Table 1). On the other hand Fst seems to perceive very ancient possible relations, like Anatolian Neolithic (AnN), Armenian Iron Age (ArmIA), European early Neolithic (EUEN), Yamnaya (Yam), Iberian Neolithic (IbN) Iberian Chalcholithic (IbCh), which appear together with their modern descendants; Armenians (Arm), Azeris (Aze), Iranians (Ira), Druze (Drz), Greeks (Gre), Sardinians (Sar), French (Fre), Scottish (Sco) etc, that are hardly detected with SHD (S9 Table). Small and negative Fst values indicate no genetic differentiation between populations, but these often couple with high P-values hence are normally not considered. However in this case the independent SHD method confirms the validity of most small and negative Fst values (S9 Table).

Table 1 lists only those populations which showed close distance values with both methods for any of the Conqueror subpopulations, as these can be considered very plausible relationships, and MDS plot of this subset is displayed on Figure 3.

**Fig. 3.**
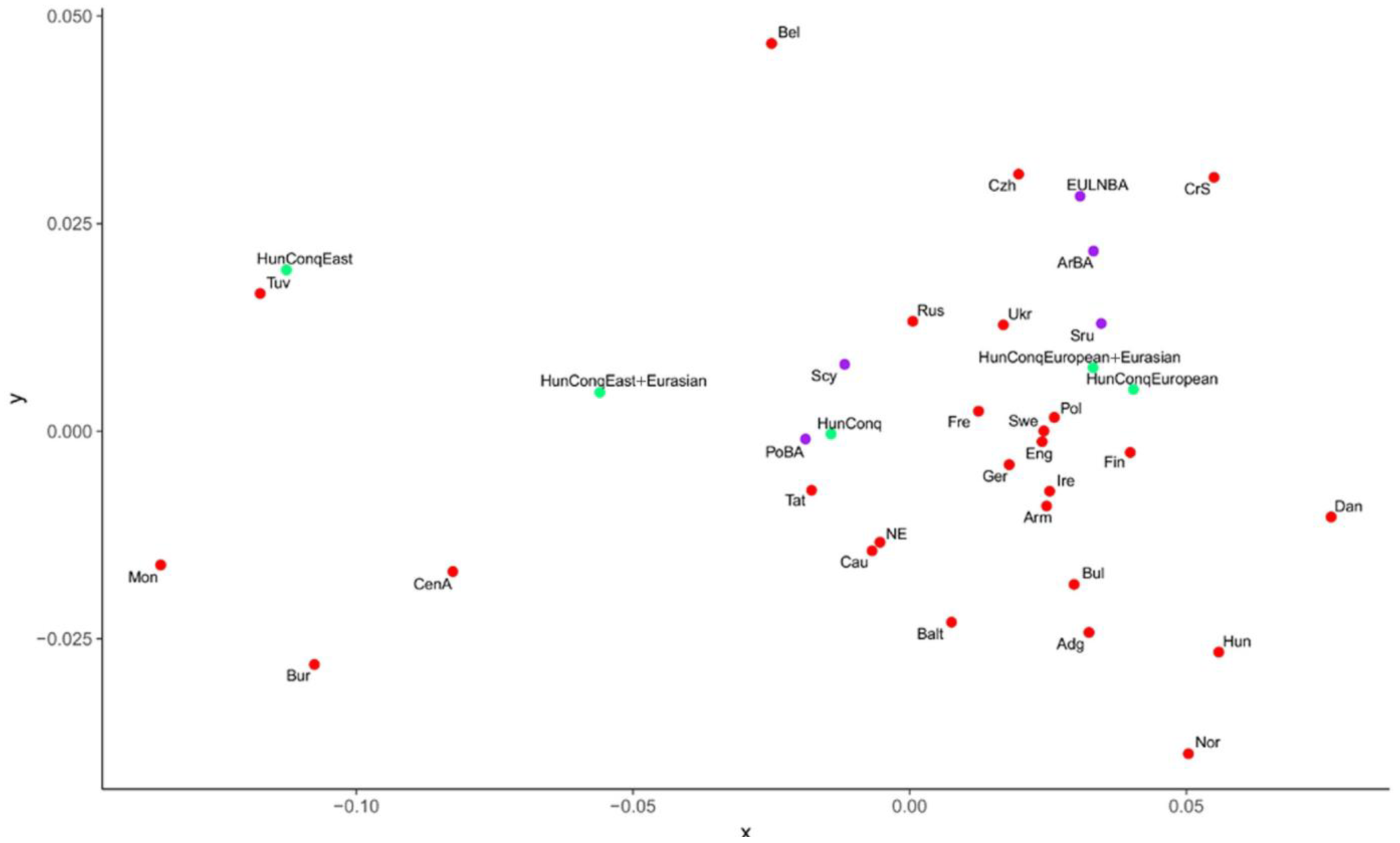
MDS plot from linearized Slatkin Fst values of S7 Table. Only populations from Table 1 were depicted, which showed close Fst and SHD distance values to the Conquerors. Abbreviations of population names are given in S4 Table.

The following pattern is discernable from Table 1; the entire Conqueror population shows nearly identical distance patterns with both methods to that of its European subset, irrespective of the presence of the 11 Eurasian lineages, while the East Eurasian component is only detected by SHD. Accordingly, the European and European+Eurasian Conqueror subpopulations map very close on Figure 3, and their closest populations are the ancient Srubnayas, modern Polish, Ukrainians, Swedish, Finnish, English, Germans, French, Irish, Armenians, and Russians. In addition Table 1 lists modern Hungarians, Danes, Croatians/Serbians, Norwegians, Belarusians, Adyghe, Bulgarians, Czechs, and Baltic populations close to the Conquerors. Of the ancient populations next to Srubnaya both methods indicate significant relation to the Poltavka-Potapovka, Armenian Bronze Age, European Scythian and European Late Neolith-Early Bronze Age populations (Table 1). From the entire Conqueror population Volga Tatars have the smallest overall distance with both methods.

Table 1 indicates that the East Eurasian Conqueror subpopulation is significantly related to modern Central Asians (Kazakhs+Uzbeks+Turkmens+Kyrgyzians) and Mongolians (including Inner Mongolians and Manchurians), which is augmented with Tuvinians and Buryats when the Eurasian component is combined with the East Eurasian one. On Figure 3 Tuvinians map closest the East Eurasian subset, while the combined subset approaches the European populations.

MITOMIX indicates that if all modern and ancient populations are considered, the Conquerors are best admixed from 26-38% modern Belarusians, 19-34% Tuvinians, 18% ancient Baltic Late Bronze Age and 13% Srubnaya populations (S10 Table). Other occurrent admix components may include Volga Tatars, Poltavka-Potapovka, Sintastha and Combed Ware populations. Thus MITOMIX principally derives Eastern Conqueror lineages from Belarusians, Tuvinians and Volga Tatars. Belarusians comprise 22% Lipka Tatars in our dataset [51], who arrived to Europe after the Conquerors’ era, but seemingly with similar Hg-s. Belarusians are best admixed from Russians, Romanians and Central Asians (S11 Table), while Tuvinians are best admixed from Central Asians and Mongolians (S12 Table).

MITOMIX derives western Conqueror lineages by augmenting the European components of above populations, with admixtures from Baltic Bronze Age (BalBA), Srubnaya and Poltavka-Potapovka populations (S10 Table). BalBA has the closest SHD distance to the Scandinavian Neolith-Bronze Age (NNBA), European Early Neolithic (EUEN) and European Scythian (Scy) groups (S8 Table), and its best admixture source is also NNBA (S13 Table).

When only ancient populations are considered as source, the best admix includes 36-44% Poltavka-Potapovka, 18-20% Baltic Bronze Age, 11-29% Combed Ware, 14-18% Sintashta and 14% Srubnaya components (S10 Table bottom), all of which are comprised of solely West Eurasian Hg-s. However ancient MITOMIX gives significantly higher SHD distances signifying that our ancient database lacks important East Eurasian components.

The most plausible interpretation of the phylogenetic and population genetic results is that most Eastern lineages were ultimately derived from Inner Asia and then migrated to Central Asia where they admixed with Eurasian lineages before moving to Europe, where they incorporated West Eurasian elements.

## Discussion

Full length mtDNA sequences facilitated a high resolution phylogenetic analysis, thus we could allocate the fairly accurate geographic origin of most individual maternal lineages. These were apparently derived from at least two rather distant source populations, one in East-Eurasia the other in Europe, raising the question as to when did the admixture happen and which ancient populations could have been the source.

### Origin of the West Eurasian maternal component

According to our data the best fitting source of the European component are the Late Bronze Age Srubnaya (Timber-grave) culture (∼1,850-1,200 BCE) and its ancestors the Potapovka (∼2,500-1,900 BCE) and Poltavka (∼2,900-2,200 BCE) cultures (Table 1). The Srubnaya was a nomadic culture on the Pontic-Caspian steppe, both their genetic composition and life style being closely related to the contemporary eastern Andronovo and Sintashta cultures together constituting the steppe Middle-Late Bronze Age (MLBA) population, which was descended from the genetically tightly clustering steppe Early-Middle Bronze Age (EMBA) Yamnaya-Afanasievo-Poltavka cultures with the addition of an European Neolithic farmer genetic layer [52], [53]. As a result, the steppe MLBA population very much resembled genetically to the European Late Neolithic/Bronze Age (EULNBA) populations [52], providing an explanation for the similarity of the Conquerors to EULNBA populations (Table 1, Figure2), the appearance of a considerable number of modern European and Northwestern European maternal lineages close to the Conquerors (S1 Figure) and the presence of European Y-chromosomal Hg-s *R1b-M269* and *I2a* in the Conquerors, reported in our previous study [13].

The Armenian Bronze Age (ArmBA) population also appears very close to the Conquerors (Table 1, Figure 3), that may be explained by the 48-58% Armenian-like Near East ancestry of the steppe EMBA populations [52], which was ultimately derived from early Iranian farmers [53]. This genetic layer may also explain the appearance of modern populations from the Caucasus region (Cau, Adg, Arm, Aze) close to the Conquerors both in population genetic (Table 1) and phylogenetic analysis, (S1 Figure). Nevertheless a more recent admixture from this region is also plausible, as all presumptive carriers of the East Eurasian lineages (Asian Scythians, Huns, Onogurs and Avars see below) contacted the Caucasus region during their westward migrations.

### Origin of the East Eurasian maternal component

Genetic and historical data refer to four major groups delivering significant East Eurasian lineages to Europe which could be connected to the Conquerors. Both anthropological [54] and genetic data [47,55] indicate that until the late Bronze Age Central Asia was populated mainly by Europid Sintashta-Andronovo people while populations with Mongoloid traits and genes were confined East of the Altai. The first Eastern Hg lineages appeared in West Siberia at the beginning of Bronze Age [56] and in the Altai at the Middle Bronze Age [57]. However on the Pontic steppe large scale appearance of Eastern Hg-s is detected just in the Iron Age when European Scythians admixed with Scytho-Siberians, giving rise to 18-26% eastern lineages in European Scythians by the 2^nd^ century BCE [49]. Western and Eastern Scythians arose independently [58], westerners were descended from the late Bronze Age Srubnaya people [59], while eastern Scythians descended from Bronze Age Andronovo people admixed with East Eurasians [49]. Our SHD and MITOMIX data corroborate these results, as European Scythians are the closest population to Srubnaya (S8 and S14 Tables). Despite the presence of Eastern lineages in European Scythians, they rather resemble to the European component of the Conquerors (Table 1) indicating that they were an unlikely source of the Eastern Conqueror lineages, but might have contributed to the European ones. There are few mitogenomic data from the Asian Scythians, but they have similar Eastern Hg composition to that of the Conquerors (Figure 4), suggesting that their descendants could have contributed to the Conqueror gene pool by later migrations.

The second major wave of East Eurasian gene flow into Central Asia, then further to Eastern Europe can be attributed to the Asian Huns (Xiongnus), whose migration from Mongolia to West through Altai and Tuva lead to a significant increase of Mongoloid anthropological components in Central Asia between the 3^rd^ century BC and 2^nd^ century AD [54,60]. During the first centuries AD Northern Xiongnus were expelled from Inner Asia and escaped westward [61]. According to some archaeologists traces of European Huns can be detected on the Pontic steppe already in the 2^nd^ century AD [62], but European Huns entered history just from the middle of the 4^th^ century as an empire. The Xiongnu origin of European Huns has been accepted by most historians [63–65], but evidences are scarce.

From Inner Asian Xiongnu remains only HVR data are available [66–69], which show the presence of predominantly Asian major Hg-s with a composition similar to the Asian subset of our Conqueror samples (Figure 4). Presence of Hg *B* in both groups is particularly informative, as it has not been detected in Scythians [49], (Figure 4) and is also missing from Western Siberian populations, but is present in modern Turkic, Mongolic, and Tungusic speaking groups of Siberia and Central Asia [70]. Phylogeographic distribution of the best East Eurasian sequence matches (S1 Figure, Figure 1 population list) center around modern Mongolia and Buryatia, which well corresponds to the territory of the ancient Xiongnu empire (Figure 7). Thus available genetic data are reconcilable with the correlation between Asian Huns and the Hungarian Conquerors.

**Fig. 4.**
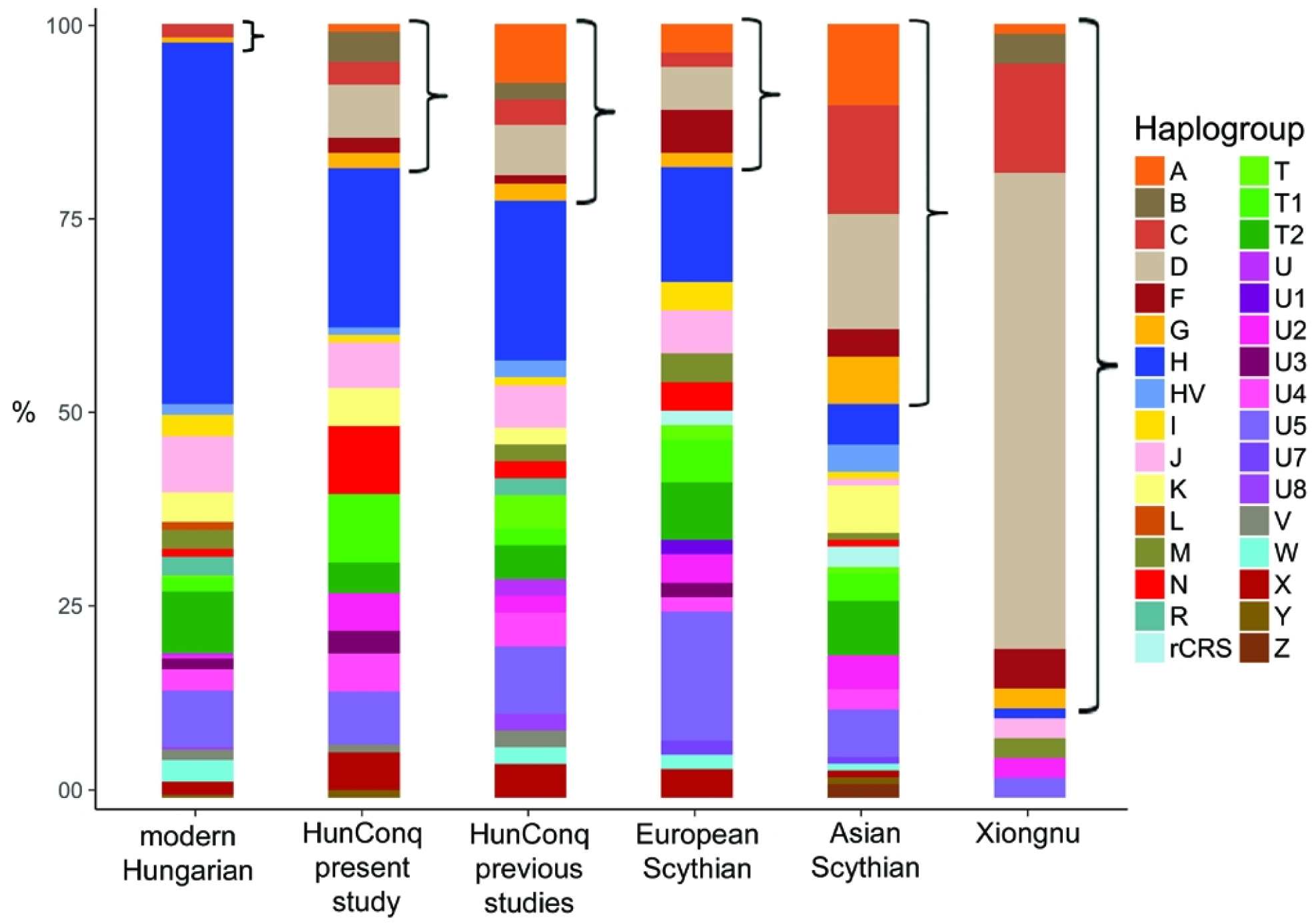
Comparison of major Hg distributions from modern and ancient populations. Asian main Hg-s are designated with brackets. Major Hg distribution of Conqueror samples from this study are very similar to that of other 91 Conquerors taken from previous studies [11,12]. Scythians and ancient Xiongnus show similar Hg composition to the bracketed Asian fraction of the Conqueror samples, but Hg B is present just in Xiongnus. Modern Hungarians have very small Asian components pointing at small contribution from the Conquerors. Of the 289 modern Hungarian mitogenomes 272 are published in [29]. Scythian Hg-s are from [48,49,55,59,71–74]. Xiongnu Hg-s are from [66–69].

There are no aDNA data from the European Huns, the only sequence from Hungary [50] belongs to the East Eurasian *D4j12* Hg which is also shared by two of our Conqueror samples (S1 Figure, Network 9) implying possible Hun-Conqueror relation.

A decade after the fall of the European Hun empire (472 AD) another grouping of Turkic tribes, the Ogurs appeared on the Pontic steppe from Central Asia. A series of sources connect the Onogurs with the later appearing Bulgars, and some to conquering Hungarians [75]. Onogur-Bulgars had been part of the Hunnic people, and after the death of Attila’s son Irnik, European Hun remains fused with the Onogurs [75]. The ensuing Avar invasion brought Onogur groups to the Carpathian Basin, others became part of the later Danube Bulgar and Volga Bulgar states. The Conquerors must have belonged to the Onogur tribal union, as the name “Hungarian” is derived from “Onogur” [5,76]. There are no aDNA data yet from Onogurs.

The succeeding group arriving from East Eurasia to the Pontic steppes in the middle of 6th century were the Avars, who established an empire in the Carpathian Basin lasting for three centuries

[77]. There are 31 HVR profiles available from the Avars [12], containing 15.3% East Eurasian lineages (*C, M6, D4c1, F1b*). However the Avars were anthropologically a mixed population with some 7.7% Mongoloid elements [78], which was virtually missing from the Conquerors (S1 Table, Figure 6). It is relevant to note that none of the Hungarian medieval sources know about Avars, presumably because they were not distinguished from the Huns [2], as many foreign medieval sources also identified Avars with the Huns [3].

Subsequent East-West migrations are connected to Göktürk, Kipchak and Mongolian groups, but these could have insignificant effect on the Conquerors as mostly arrived after the 10^th^ century, moreover most Turkic loanwords in Hungarian originate from West Old Turkic [79], the Oghur Turkic branch associated with previous Turkic speaking groups as Onogurs, Bulgars, Khazars and maybe the Avars.

**Fig. 6.**
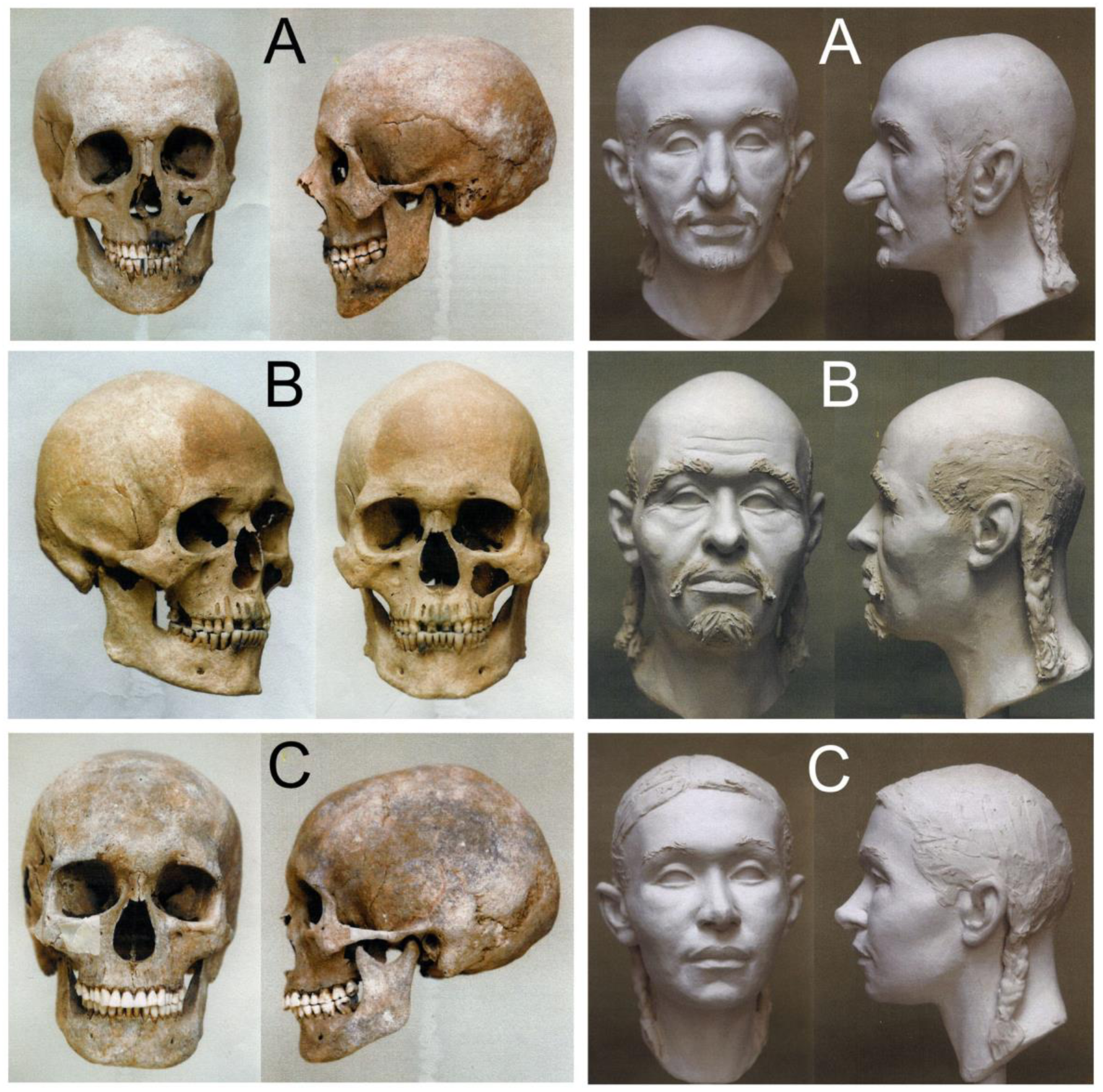
Skulls and sculpting craniofacial reconstructions of Hungarian Conqueror individuals. **A**: Karos2/52 mature aged leader with Europid anthropological features. **B**: Karos2/60 senile aged man with Europo-Mongoloid features. **C**: Karos2/47 adult woman with Europo-Mongoloid features.

### Relation to Volga Tatars

From all recent and archaic populations tested the Volga Tatars show the smallest genetic distance from the entire Conqueror population with both methods (Table 1, Figure 3), which can be explained either by direct genetic relation or analogous population history. The history of Volga Tatars suggests that this similarity accumulated from multiple historical episodes. Volga Tatars (and their Bashkir neighbors) incorporate three main ethnic components; the Volga Bulgars, which arrived in the 8^th^ century, and intermingled with local Scythian and Finno-Ugric populations, then in the 13^th^ century Kipchak Tatars of the Golden Horde brought a final Central-Inner Asian genetic layer and their language to the region [80]. Thus both the East and West Eurasian component of the Volga Tatars derives from multiple sources and much of their East Eurasian component arrived to this region after the Conquerors’ era, but from the same Central-Inner Asian source. This is also manifested by the considerable number of other Tatar groups (Belarusian, Altai, Kyrgyz) appearing close to the Conquerors on the phylogenetic trees (S1 Figure, Networks 9, 12, 23, 50). It is remarkable that according to MITOMIX the Conquerors seem to provide a dominant (26-41%) component of the Volga Tatars besides the Russians and Finno-Ugric Yugra people (S15 Table), indicating that rather Volga Tatars harbor a “Conqueror like” genetic component than the opposite. Historically this component may be linked to the Volga Bulgars, who were one of the few groups having the same partial horse burial [81] and symbolic trepanation customs [82,83] as the Conquerors.

There are limited mitogenomic data available from the neighboring Bashkir population, but they have a similar history [84] and genetic composition to Volga Tatars, moreover also contain low frequency *B*, *Y* and *N1a1a1a1a* Hg-s characteristic to the Conquerors and missing from Volga Tatars [85] indicating an even closer genetic link obtained from similar sources. So a direct genetic relation of the Conquerors to Onogur-Bulgar ancestors of these groups is very feasible.

### Finno-Ugric relations

Surprisingly we did not find significant genetic relations to Finno-Ugric groups. Though population genetic analysis indicates some connection of the Conqueror European component to modern Finnish (Fin) and Baltic people (Balt=Latvians+ Lithuanians+Estonians) (Table 1), but no relation to Saamis (Sam), Mansis and Kanthys (Yug) (Table 1, S9 Table). The Baltic relation of the European component seems to appear already in the Baltic Bronze Age (BalBA, 1000-230 BCE), [86] measured with the SHD method (Table 1, sequence data are not yet available). BalBA genomes cluster with modern Lithuanians and Estonians, and lack Eastern mtDNA Hg-s and Y-chromosomal haplogroup *N-tat*, which is typical for Uralic speaking groups, thus Estonians must have received their East Asian-Siberian components after the BalBA period, from a different source [86]. According to our data BalBA is best admixed from the closely related Scandinavian Neolith-Bronze Age (NNBA), Afanasevo and European Neolithic populations (S13 Table), so it is unlikely connected to Finno-Ugric groups. As only 7 Estonian mitogenomes are available, they were grouped with other modern Baltic populations (Balt), so the similarity of these to the Conquerors probably derives from BalBA heritage. The connection to modern Finnish population can also be explained from the BalBA and steppe MLBA component which is present in modern Scandinavians, as Finnish sequence matches regularly appear together with Danish ones on our phylogenetic trees (S1 Figure, Networks 14, 15, 19, 25, 27, 30, 35, 40, 42, 43, 49, 52, 56).

Moreover, *Y*, *B* and *N1a1a1a1a* Hg-s have not been detected in Finno-Ugric populations [87], [88], [89], [70] [85] indicating that the East Eurasian component of the Conquerors and Finno-Ugric people are probably not directly related. The same inference can be drawn from phylogenetic data, as only two Mansi samples appeared in our phylogenetic trees on the side branches (S1 Figure, Networks 1, 4) suggesting that ancestors of the Mansis separated from Asian ancestors of the Conquerors a long time ago. This inference is also supported by genomic Admixture analysis of Siberian and Northeastern European populations [90], which revealed that Mansis have very ancient North Eurasian ancestry, who in addition received a significant (43%) Eastern Siberian genetic component approximately 5-7 thousand years ago from ancestors of modern Even and Evenki people. Most likely the same explanation applies to the Y-chromosome N-Tat marker which originated from China [91], [92] and its subclades are widespread between various language groups of North Asia and Eastern Europe [93].

It must be emphasized that Finno-Ugric groups are underrepresented in our population database, as we have no mitogenomic data from Komis, Maris, Mordvins and Udmurts and only limited samples from Mansis, Kanthys, Saamis and Estonians, thus appearance of Finno-Ugric matches from a more representative dataset cannot be excluded. The Poltavka, Potapovka, Scythian and Volga Tatar relations indicate that the Samara district might have contributed to the European Conqueror component multiple times, and the Upper Volga region was also a probable source of Uralic people between 5300-1700 BCE [94]. Thus our data imply that incidental Finno-Ugric link is rather expected in the European component if any.

### Genetic relation of different Conqueror cemeteries

Archaeologist presume that the rich 10th century cemeteries of Karos and Kenézlő comprise the Conqueror military elite, raising the question as to what extent can our findings be generalized to the entire Conqueror population. Our fragmentary data from other cemeteries indicate the presence of the same Eastern and Western genetic components (S1 Figure, Networks 3, 4, 12, 36), moreover [12] and [11] reported 91 other Conqueror HVR haplotypes from 24 cemeteries, which show very similar major Hg distribution to our samples (Figure 4), with even larger proportion of Asian major Hg components. Thus our conclusions probably apply to the entire Conqueror population, but definitely to the 10^th^ century immigrant military elite characterized with partial horse burials, though further mitogenomic and genomic data are required for the accurate answer.

We have determined the maternal lineage of the majority of samples from the three neighboring Karos cemeteries, and found likely maternal relatives with identical mtDNA genomes within cemeteries allocated into the same circles on the phylogenetic trees, on S1 Figure 1 (summarized in S2 Table1), but surprisingly no identical haplotype was found between the three Karos cemeteries. The only exceptions are the two chiefs in Karos2 and 3, who had identical *X2f* maternal haplotypes and *I2a1* Y chromosomal haplotypes (data not shown), so were probably brothers. This indicates that these neighboring communities did not intermarry probably because of different group-identity. Furthermore the East Eurasian haplogroup lineages from the three Karos cemeteries indicate a discernible structuring (S2 Table2); the Karos3 cemetery has a definite South-East Chinese affinity, the Karos1 a North-East Siberian affinity, while the Karos2 lineages are widely distributed from East to Central Asia. In contrast, despite the low number of samples analyzed from other Conqueror cemeteries we detected potential relatives with identical mtDNA genomes between distant cemeteries (S2 Table1). This suggests that individual tribes might have been split and fragments of different tribes settled together upon the conquest.

### Relation of Conquerors to modern Hungarians

Modern Hungarians are genetically very similar to their European neighbors [95] nevertheless they contain some 3-5% East Eurasian components traceable with uniparental markers [29,96,97]. Genome wide SNP data also detected the presence of 4% East Asian component in modern Hungarians [98] with an approximate time of admixture dated to the first millennium AD, corresponding to the invasions of Huns, Onogur-Bulgars, Avars and Hungarian Conquerors from the Asian steppes. These findings are completely in line with our results, as the Hun, Onogur, Avar and Conqueror East Eurasian genetic components seem inseparable indicating that they represent similar source populations ultimately derived from Xiongnus admixed with descendants of Asian Scythians.

Thus genetic heritage of the Conquerors definitely persists in modern Hungarians, but both the ratio of East Eurasian components (30:3%) and MITOMIX data [29] indicate that they contributed to less than 1/10^th^ of recent Hungarian gene pool. This dilution could have started at the time of the conquer, as contemporary local population size in the Carpathian Basin was estimated larger than that of the Conquerors [99,100]. Anthropological data also have the same implication, as the Conquerors differed from the subsequent Árpádian Age population, which was more similar to preconquest Avar Age populations [101,102]. According to early anthropological studies people of the Avar and Conquest age Carpathian Basin were very heterogeneous and immigrants arrived in several phases between the 5^th^ and 9^th^ centuries [103], which in our view admixed with the autochthonous population, of which genetic data are still barely available between the Bronze Age and Conquest period.

The large genetic diversity of the Conquerors which seemingly assembled from multiple ethnic sources and their relative low proportion, having no lasting effect on Hungarian ethnogenesis, raises doubts about the Conqueror origin of the Hungarian language. Even if our samples represent mainly the Conqueror elite, the “elite dominance” linguistic hypothesis seems inconsistent when it presumes that the same Turkic elite was first readily assimilated linguistically by Finno-Ugric groups, and then it assimilated locals of the Carpathian Basin. Turkic character of the Conqueror elit is indicated by their “Turk” denomination in contemporary sources as well as Turkic tribal names and person names of tribe leaders of the conquest-period [104]. Above data infer that preconquest presence of the language in the Carpathian Basin, is an equally grounded hypothesis, as had been proposed by several scientists (a summary in English is given in [105]), which is also hinted by a recently detected genomic admixture between Mansis and a Middle Neolithic (5000 BCE) individual from the Carpathian Basin [90].

## Conclusion

The large diversity of Hg-s detected in the Conquerors reflects a quite complex genetic history, which was summarized from our data on Figure 7. Their uniform archaeological findings and predominantly Europid anthropological features (Figure 6) indicate a long lasting admixture on the Pontic steppe, thus their final composition was likely formed there during the last centuries prior to the conquest. Members of each groups bringing eastern Hg lineages to Europe could have originated from Xiongnu and Asian Scythian foremothers. On the Pontic steppes Asian nomads assimilated with descendants of the Srubnayas and this mixed population could have been the basis of many medieval Pontic nomadic groups, including Conquerors. Their ancestors were certainly part of the European Hun Empire, the succeeding Avar and Bulgar empires, and when they came into power very probably incorporated European Hun remains, as recognized previously [106]. Our genetic data support the Hun-Conqueror connection which could have been the basis of the historical-cultural Hungarian Hun tradition [3]. As a consequence our data provide indirect genetic evidence for the thus far debated Xiongnu origin of European Huns as well. Genetic relation of the Conquerors to medieval Onogur-Bulgars warrants further studies, as they are linked by archaeological, anthropological and historical data as well as our population genetic indications.

Our conclusions are well supported by anthropological studies, which found analogies of the lower class Conqueror individuals on the Eastern European steppes, but parallels of the upper warrior class were mainly found at the fringes of the Xiongnu empire, in South Siberia and South-Central Asia [107]. Finally our data indicate that all potential ancestors of the Conquerors were steppe nomadic people, which is in full agreement with their archaeological legacy.

**Fig. 7.**
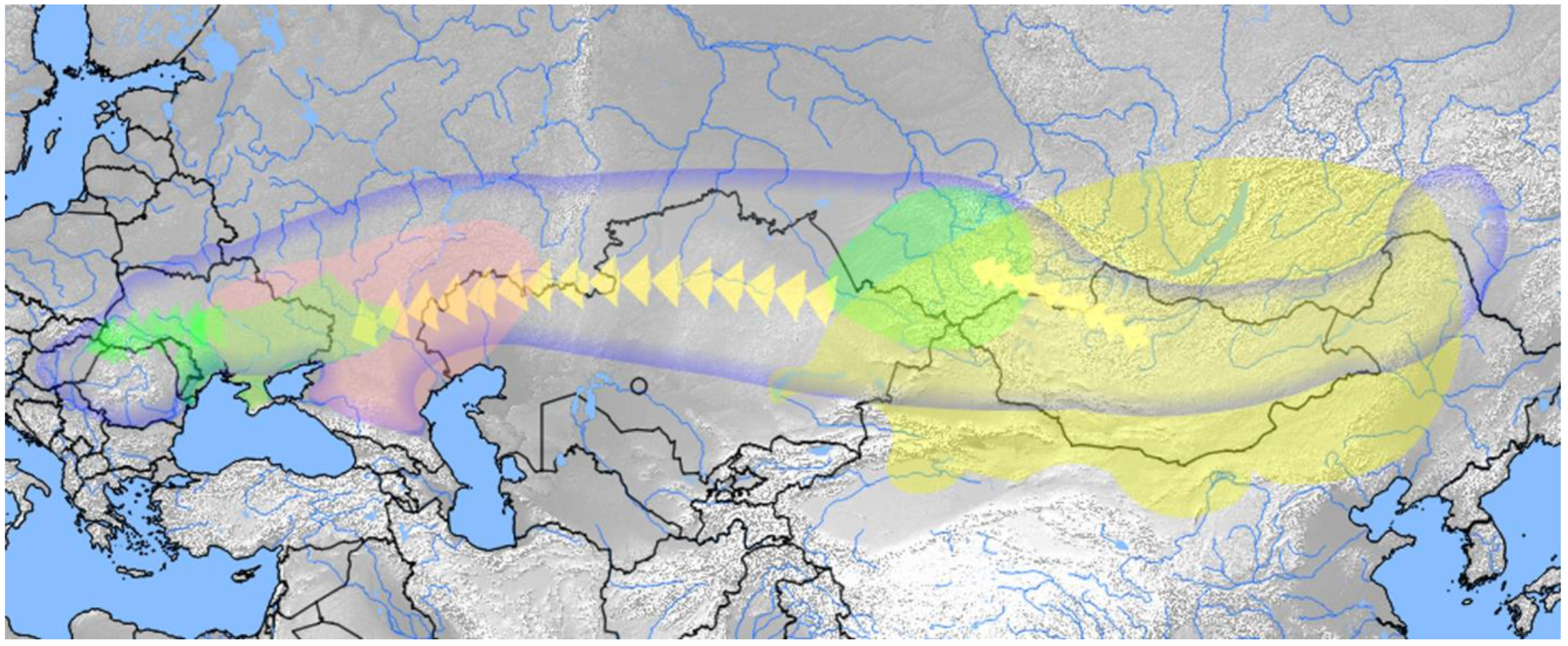
Hypothetic origin and migration route of different components of the Hungarian Conquerors. Bluish line frames the Eurasian steppe zone, within which all presumptive ancestors of the Conquerors were found. Yellow area designates the Xiongnu Empire at its zenith from which area the East Eurasian lineages originated. Phylogeographical distribution of modern East Eurasian sequence matches (Fig. 1) well correspond to this territory, especially considering that Yakuts, Evenks and Evens lived more south in the past [108], and European Tatars also originated from this area. Regions where Asian and European Scythian remains were found are labeled green, pink is the presumptive range of the Srubnaya culture. Migrants of Xiongnu origin most likely incorporated descendants of these groups. The map was created using QGIS 2.18.4[109].

## Supplementary Material

**Supplementary text:** Archaeological background

**S1 Figure:** Median-Joining Networks (1-58) for mtDNA sequences of the 102 Hungarian Conquerors.

**S1Table:** Description of samples, including anthropological and archaeological details.

**S2 Table:** List of samples with Identical mtDNA sequences, Distribution of the East Eurasian Hg-s in the three Karos cemeteries.

**S3 Table:** Details of NGS data for each samples.

**S4 Table:** Summary of the modern (top) and ancient (below) population database with abbreviations used in this study.

**S5 Table:** Population database assembled from NCBI and ENA, showning mitogenome Hg-s described from each population together with their frequencies in the given population.

**S6 Table:** Conqueror subpopulations considered in population genetic analysis.

**S7 Table**: Pairwise Fst (top) and linearized Slatkin Fst (below) matrix of population distances between all combinations of modern and ancient populations.

**S8 Table:** Pair-wise Shared Haplogroup Distance (SHD) values measured between all combinations of modern and ancient populations.

**S9 Table.** Comparison of population genetic distance values measured with two different methods (Fst and SHD) between Hungarian Conqueror subpopulations and all ancient (arch) and modern (rec) Eurasian populations.

**S10-15 Tables**: MITOMIX results for the entire Conqueror population and for the source populations of the best MITOMIX: modern Belarus, Tuvan, Baltic Bronze Age, Srubnaya and Volga Tatars populations from available population Hg frequency data.

## Availability of data and materials

The raw sequence data of the 102 samples were deposited to the European Nucleotide Archive (http://www.ebi.ac.uk/ena) under accession number PRJEB21279.

## Author contributions

T.T. and E.N. designed research; E.N., T.T., K.K. and K.M. performed archaic DNA experiments, NGS library preparation and enrichment; P.B. and I.N. evaluated library quality and performed NGS sequencing; Z.M. implemented bioinformatics analysis; Z.M., E.N., T.T. and T.K. analyzed data; E.F. assembled archaeological background; Á.K. made craniofacial reconstructions; E.F., I.P., Á.K. and Gy.P. provided samples and anthropological data; I.R. provided financial and scientific support; A.Z. facilitated initial phases of NGS methods; T.T. wrote the manuscript, and all authors reviewed and approved the final version. E.N. and M.Z. contributed equally to this study.

## Acknowledgement

We would like to thank Alissa Mittnik, Mark Stoneking, Leyla Dzhansugurova and Eppie R. Jones for providing unpublished sequences. We also thank Kornél Bakay, Zoltán Kristóf, László Révész, Pál Sümegi and Balázs Tihanyi for their useful advices in topics of archaeology and history. Erika Molnár, and András Bíró helped us with anthropological materials and background, while Zoltán Juhász called our attention to folk music relations. This research was supported by grant no. GF/JSZF/814/9/2015 to I.R, and in part by grant NKFI-6 no. K-124350 to TT.

